# ILC3 inhibits Osteoclastogenesis and ameliorates Inflammatory Bone Loss in Post-menopausal Osteoporosis

**DOI:** 10.1101/2025.09.20.677486

**Authors:** Asha Bhardwaj, Leena Sapra, Simran Preet Kaur, Tamanna Sharma, Chaman Saini, Pradyumna K. Mishra, Rupesh K. Srivastava

**Affiliations:** Translational Immunology, Osteoimmunology & Immunoporosis Lab (TIOIL), An ICMR Collaborating Centre for Excellence on Bone Health, Department of Biotechnology, All India Institute of Medical Sciences (AIIMS), New Delhi-110029, India; Department of Molecular Biology, ICMR-National Institute for Research in Environmental Health, Bhopal, MP, 462001, India

**Keywords:** Innate lymphoid cell (ILC)-3, Bone, Post-menopausal osteoporosis, short chain fatty acids (SCFAs), T-bet^+^ ILC3, Dysbiosis

## Abstract

Although the significance of T lymphocytes in maintaining bone homeostasis is well established, the role of innate lymphoid cells (ILCs), the innate counterparts of T cells, in maintaining bone homeostasis is uncertain. In this study, we examined, for the first time, the anti-osteoclastogenic property of ILC3 *in vitro*. We observed that ILC3 inhibits RANKL-mediated osteoclastogenesis in a cell ratio-dependent manner. We further employed an ovariectomized (ovx) mouse model, which mimics postmenopausal osteoporosis (PMO), to investigate the role of ILC3 in inflammatory bone loss. Notably, our *in vivo* data unequivocally validate that the ovx mice have a markedly lower frequency of ILC3 in the bone marrow (BM). Furthermore, the temporal-kinetic analysis revealed that alterations in ILC3 cell dynamics and function in the BM are associated with the development and progression of inflammatory bone loss in PMO. Additionally, our *in vivo* results demonstrate that dysbiosis in PMO drives significant alterations in the frequency of ILC3 subsets by promoting the expansion of IL-17-producing Nkp46^−^ ILC3 and inhibiting the development of IL-22-producing Nkp46*^+^* ILC3. Moreover, T-bet^+^ ILC3s are significantly increased in ovariectomized (ovx) mice. Altogether, the present study for the first time, reports the critical role of dysregulated ILC3 in the progression of PMO and offers a novel immunotherapeutic approach targeting ILC3 for treating and managing PMO.

**Graphical abstract:** PMO is associated with leaky gut and dysbiosis, which leads to a deficiency of SCFAs in the ovx mice. Dysbiosis in ovx mice disturbs the homeostatic balance of the ILC3 with an increase in IL-17-producing ILC3 and a decrease in IL-22-producing ILC3. Further Tbet expression is enhanced in the ILC3 of the ovx mice, which is associated with disease pathogenesis. Additionally, IL-2 expression is decreased in the ILC3 of the ovx mice. Decreased expression of ILC3 migratory molecules reduced the migration of ILC3 in bone marrow. ILC3 inhibits osteoclast differentiation. However, in the case of ovx mice, ILC3 are compromised and promote osteoclastogenesis.

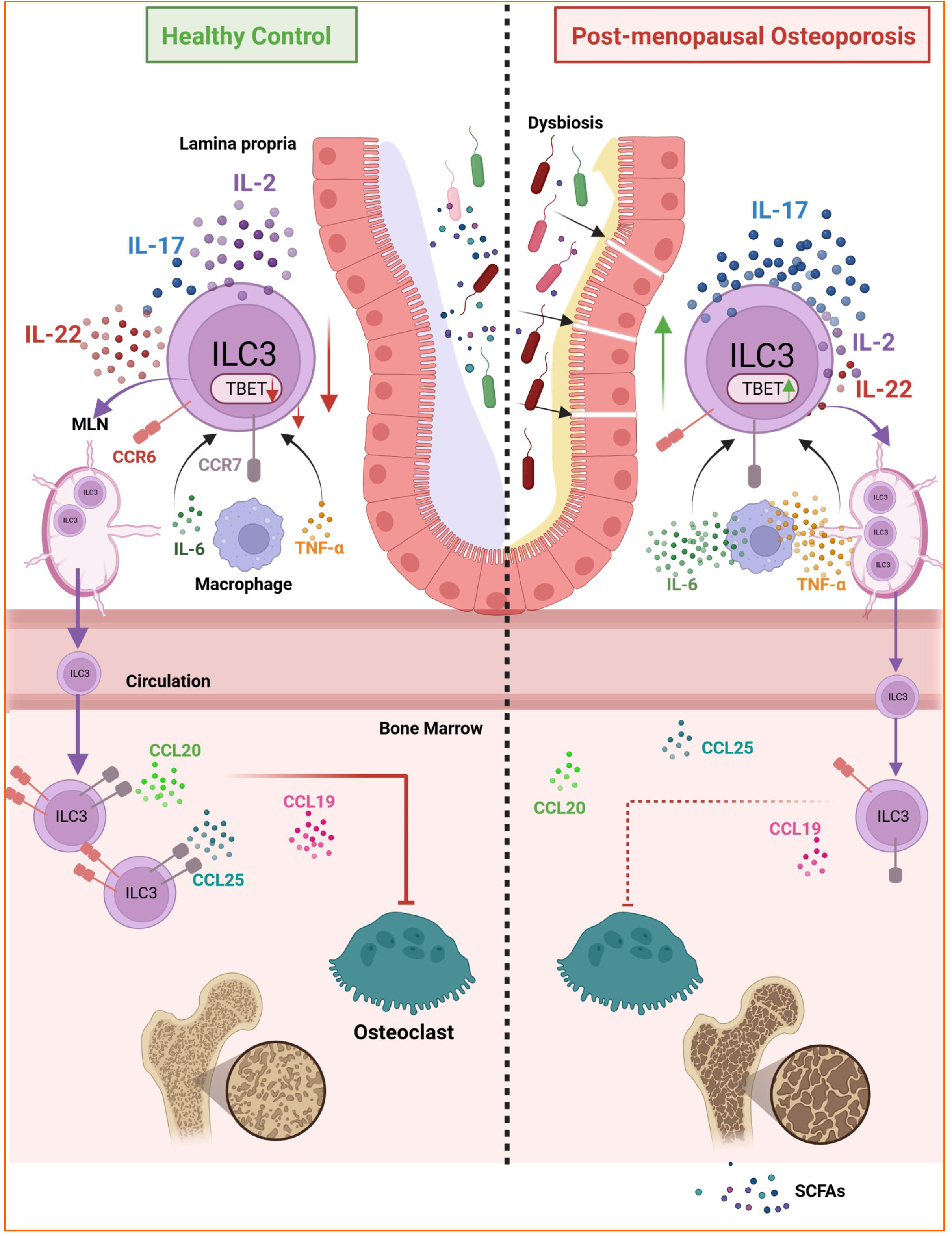

## 1. Introduction

Osteoporosis is a systemic skeletal disorder characterized by decreased bone mineral density (BMD), which raises the risk of bone fractures. Osteoporotic fractures greatly reduce quality of life and increase morbidity, mortality, and disability. Since bone loss often occurs without symptoms, osteoporosis is often called a “silent disease” that can affect both sexes and worsens with age. The pathophysiology of osteoporosis is complex; although traditional views focus on endocrine pathways, recent studies reveal that the immune system also strongly influences bone (Guder et al., 2020). Over the past 20 years, numerous researchers have investigated the relationships between the immune system and bone cells. Along with their key role in immune responses, T cells and B cells can also influence bone remodeling. Evidence shows that Th1, Th2, and Treg cells inhibit osteoclast formation through their cytokines, whereas Th17 and Th9 cells promote osteoclast development (Andreev et al., 2022; Dar, Azam, et al., 2018; Sapra et al., 2024; Srivastava & Sapra, 2022a). Researchers have uncovered a complex relationship between B cells and bone cells. It has been demonstrated that B cells in the bone marrow of estrogen-deficient postmenopausal women release receptor activator of NF-ĸB ligand (RANKL) (Onal et al., 2012), providing a mechanism for B cells’ role in estrogen deficiency-related bone loss. In an *in vitro* human model of osteoclastogenesis, peripheral blood B lymphocytes inhibit osteoclast maturation by secreting transforming growth factor (TGF)-β (Weitzmann et al., 2000). Our lab was the first to identify the role of regulatory B cells (Bregs) in suppressing osteoclastogenesis *in vitro*, a process mediated by IL-10 production (Sapra, Bhardwaj, et al., 2021; Sapra et al., 2025). With greater understanding of the complex relationship between bone and the immune system in recent decades, our group introduced the novel concept of “immunoporosis,” which highlights the expanding role of immune cells in osteoporosis (Pol et al., 2023; Sapra, Azam, et al., 2021; Srivastava et al., 2018; Srivastava & Sapra, 2022).

Not only the adaptive immune system but also the innate immune system is intricately involved in bone physiology and pathologies. Innate immune system cells such as macrophages and dendritic cells are linked to managing bone homeostasis, which involves eliminating apoptotic cells, executing autophagy, and resolving inflammation caused by cellular injury (Charles & Nakamura, 2014; Xiang & Gilkes, 2019). Innate lymphoid cells (ILCs) are a group of newly discovered innate immune cells that have become crucial effectors of innate immunity in various physiological functions. These cells are mirror images of T lymphocytes. Therefore, ILCs respond quickly to infections and other danger signals like innate immune cells, while secreting a cytokine profile similar to T cells despite lacking T cell receptors (Vivier et al., 2018). While the roles of various adaptive and innate immune cells in regulating bone cells are known, the role of the newly identified ILCs remains unclear. One study has shown that ILC2 suppresses osteoclast formation by releasing cytokines such as IL-4 and IL-13, playing an important role in regulating bone homeostasis (Omata et al., 2018). Consequently, because ILCs are mirror images of T cells, they might also have the potential to influence bone health.

In this context, type 3 ILCs (ILC3), which are typically involved in the development of secondary lymphoid organs during embryonic stages (Savage et al., 2017) and are also known to support gut integrity and fight infections (Serafini et al., 2022), are particularly intriguing. They release cytokines related to type 3 immune responses, such as IL-17 and IL-22 (Van Maele et al., 2010), some of which are also recognized as regulatory cytokines in bone homeostasis (Bugaut & Aractingi, 2021a; Triggianese et al., 2016). Notably, type 3 immune responses have been shown to possess regulatory, anti-inflammatory properties in arthritis and spondylitis, diseases strongly linked to disrupted bone homeostasis (Mauro et al., 2019). Although these findings suggest an indirect influence of ILC3 on bone through inflammation, a direct effect of ILC3 on bone, especially on bone-resorbing osteoclasts, has not been confirmed. To date, no established communication exists between ILC3 and bone cells (such as osteoclasts). In this study, we focus on examining the role of ILC3 in osteoclastogenesis and its influence on bone health under estrogen-deficient postmenopausal osteoporotic conditions. Overall, our data highlight the vital role of ILC3 in preventing inflammatory bone loss in PMO. We suggest that targeting ILC3 through immunotherapy could be a promising strategy for treating and managing PMO.

## 2.0 Materials and Methods

### 2.1 Animals

C57BL/6J female mice, aged 8-10 weeks, were housed under specific pathogen-free conditions at the Central Animal Facility of the All India Institute of Medical Sciences (AIIMS), New Delhi, India, and all *in vivo* and *in vitro* experiments were performed. For the *in vivo* experiment, the study included two groups: sham (control) and ovariectomized (ovx), with four mice in each group. Both groups were kept in separate cages with 12-hour light and dark cycles. Food and water were available ad libitum. At the end of 45 days, mice were euthanized by carbon dioxide asphyxiation, and blood, bones, mesenteric lymph nodes (MLN), and small intestine (SI) were collected for further analysis. The procedure was conducted in accordance with established principles and guidelines, following approval from the Institutional Animal Ethics Committee of AIIMS, New Delhi, India (383/IAEC-1/2022).

### 2.2 Antibodies and reagents

For flow cytometry, the following antibodies and kits were procured: CD3ε (BD, 553059), CD4 (BD, 553044), CD19 (BD, 553784), F4/80 (eBioscience, 13-4801-82), NK1.1 (BD, 553163), FCεRI (eBioscience, 13-5898-82), Gr-1 (BD, 553124), CD5 (BD, 553018), CD11c (BD, 553800), Ter119 (BD, 553672), streptavidin microbeads (BD, 557812), APC streptavidin (BD, 554067), PE-Cy7 anti-mouse-CD3 (BioLegend, 100220), APC anti-mouse-IL-22 (eBioscience, 17-7222-80), BV786 anti-mouse-IL-17 (BioLegend, 506928), PE anti-mouse-RORγT (eBioscience, 12-6988-82), BV711 anti-mouse-Tbet (BioLegend, 644820), Foxp3/transcription factor staining buffer (eBioscience, 0-5523-00), and RBC lysis buffer (eBioscience, 00-4300-54). Phorbol myristate acetate (PMA) (P1585), ionomycin (10634), and acid phosphatase leukocyte (TRAP) kit (387A) were procured from Sigma-Aldrich (USA). Monensin (420701) was obtained from BioLegend. The Mouse IL-17 (M1700) ELISA kit was purchased from Elabscience. Macrophage colony-stimulating factor (MCSF) (300-25) and sRANKL (310-01) were procured from PeproTech (USA). α-Minimal essential medium (MEM) and Roswell Park Memorial Institute (RPMI)-1640 medium was obtained from Gibco (Thermo Fisher Scientific, USA).

### 2.3 Purification and activation of ILC3s

A single-cell suspension of spleen was prepared, and after RBC lysis, the resulting cells were incubated with a biotinylated mouse ILCs enrichment cocktail (Lin-(CD3, CD19, F4/80, NK1.1, FCεRI, Gr-1, CD5, CD11c, TER119) (BD, USA) for 30 minutes at 4°C. Labeled cells were then washed with 1X PBS and incubated with streptavidin particles plus DM for 30 minutes at 4°C. Next, the cells underwent magnetic separation using a cell separation magnet, and a negative fraction containing the resulting ILCs was collected and assessed for purity (>90%) by flow cytometry. The purified ILCs were then cultured in 96-well plates (50,000 cells) with and without ILC3-surviving cytokines (IL-2 and IL-7, 10 µg/ml) and stimulating cytokines (IL-23 and IL-1β, 50 µg/ml) for 24 hours at 37°C in a humidified 5% CO[incubator. At the end of the experiment, the ILC3 population was analyzed by flow cytometry.

### 2.4 Osteoclast differentiation and tartrate-resistant acid phosphatase (TRAP) staining

Mouse bone marrow cells (BMCs) were harvested from the femur and tibiae of 8–12-week-old female C57BL/6 mice, and RBC lysis was performed using 1X RBC lysis buffer. Cells were cultured overnight in a T25 flask in α-MEM media supplemented with 10% heat-inactivated fetal bovine serum (FBS) and MCSF (35 ng/ml). The next day, non-adherent cells were collected and seeded in a 96-well plate (50,000 per well) in osteoclastogenic medium supplemented with MCSF (30 ng/ml) and RANKL (60 ng/ml), with or without IL-2 at various concentrations (1, 2, 5, 10, 20, 50, and 100 ng/ml). On day 3, half of the media was replaced with fresh complete α-MEM media containing the cytokines. To assess the formation of mature multinucleated osteoclasts, an osteoclastogenic assay using TRAP staining was performed. At the end of incubation, cells were washed twice with 1× PBS, fixed with a solution of citrate, acetone, and 3.7% formaldehyde for 10 minutes at 37°C. After fixation, cells were stained for TRAP using solutions A and B as specified by the manufacturer, incubated at 37°C in the dark for 10–15 minutes. Multinucleated TRAP-positive cells containing three or more nuclei were considered mature osteoclasts. These TRAP-positive multinucleated cells were counted and imaged using an inverted microscope (ECLIPSE, TS100, Nikon). The area of TRAP-positive cells was quantified with ImageJ software (NIH, USA).

### 2.5 Co-culture of ILC3 with BMCs for Osteoclastogenesis

Mouse BMCs were harvested from the femur and tibiae of 8–12-week-old female C57BL/6 mice, and RBC lysis was performed using 1X RBC lysis buffer. Cells were cultured overnight in a T25 flask in α-MEM media supplemented with 10% heat-inactivated FBS and MCSF (35 ng/ml). The next day, non-adherent cells were collected and seeded in a 96-well plate (50,000 per well) in osteoclastogenic medium supplemented with MCSF (30 ng/ml) and RANKL (60 ng/ml), with or without differentiated ILC3s at ratios of 1:1 and 1:5 for 4 days. The media was replenished on the third day by removing half and replacing it with fresh media containing the supplements. To evaluate the formation of mature multinucleated osteoclasts, TRAP staining was performed.

### 2.6 Flow Cytometry

Cells were harvested from BM, MLN, and SI and stained using antibodies specific to ILC3. For surface markers, cells were incubated for 30 minutes with biotin-labeled antibodies for lineage markers on ice. After washing, the cells were stained for CD45 and APC-streptavidin. Cells were then fixed, permeabilized, and incubated with anti-RORγT-PE for 45 minutes in the dark on ice. After washing and adding FACS buffer, cells were acquired on a flow cytometer (Symphony). FlowJo-10 (TreeStar, USA) software was used to analyze the samples. For analysis of intracellular IL-17 and IL-22 cytokines by ILC3s, isolated cells from BM, MLN, and SI were resuspended in RPMI-1640 complete media and activated with PMA (50 ng/mL, Sigma Aldrich), Ionomycin (500 ng/mL, Sigma Aldrich), and protein transport inhibitor Monensin for 5 hours. After surface staining with anti-CD3-PECy7, cells were fixed and permeabilized. They were then stained with anti-RORγT-PE, anti-IL-17-BV786, and anti-IL-22-APC and incubated for 45 minutes in the dark on ice. After washing and adding FACS buffer, cells were acquired on a flow cytometer (Symphony).

### 2.7 Post-menopausal osteoporotic mice model (PMO)

In the ovx group, mice underwent bilateral ovariectomy after being anesthetized via intraperitoneal injection. A nearly 1 cm incision was made on the ventral side to expose the internal organs. The uterine horn was then used to locate both ovaries. Both fallopian tubes were tied with sterile thread, and the ovaries were carefully removed with scissors. The peritoneal layer was sutured, followed by suturing the skin with sterile thread. Post-operative care was provided for one week.

### 2.8 Cytokine analysis by Enzyme-Linked Immunosorbent Assay (ELISA)

Blood was collected via the retro-orbital route and left to settle for 45 minutes at room temperature, then centrifuged at 4000 rpm for 20 minutes. The serum was collected and stored at −80°C for cytokine analysis. ELISA was performed to quantitatively measure the osteoclastogenic cytokine, namely IL-17, in the blood serum.

### 2.9 Scanning Electron Microscopy (SEM)

As previously mentioned, SEM was used to examine the cortical region of femoral bones (Sapra et al., 2022). Bone samples were kept in 1% Triton-X-100 for two to three days before being transferred to 1% PBS buffer for the duration of the final study. After preparing the bone slices, the samples were dried under an incandescent lamp before sputter-coating. Then, using a Leo 435 VP microscope with a 35-mm photographic system, the bones were scanned. Digital photos of SEM images taken at a magnification of 100 were used to capture the finest cortical region.

### 2.10 Micro-computed Tomography (µCT)

Samples were kept in 4% PFA for 3-4 days before being transferred to 1X PBS for further analysis. μCT of the lumbar vertebrae (LV)-5, femur, and tibia bones was performed using a SkyScan 1176 scanner (SkyScan). Scanning was conducted at 100 kV, 100 mA, with a 0.5 mm aluminum filter and an exposure time of 590 ms. A total of 1800 projections were collected at a resolution of 6.93 mm per pixel. The reconstruction process was carried out using NRecon software. CT Analyzer software (version 1.02; SkyScan) was used for morphometric quantification of trabecular and cortical bone indices. The Batman software was utilized for the processing of 3D and 2D analyses of LV-5, femur, and tibia cortical bones.

### 2.11 Histology

For histological analysis, SI was collected from both the sham and ovx groups. Tissues were fixed in 10% formaldehyde solution at room temperature overnight for histological staining. After paraffin sectioning, slides were stained with hematoxylin and eosin. Slides were hydrated after completely removing the wax, followed by nuclear staining with hematoxylin. Excess stain was rinsed off under tap water. Counterstaining with eosin was then performed, followed by dehydration and mounting. Tissue histological changes were examined using a microscope equipped with a digital camera.

### 2.12 qPCR

Gene expression was measured using RT-PCR (Applied Biosystems, Quantstudio™-5, USA). Duplicate samples of cDNA from each group were amplified with customized primers, including RANKL, OPG, CCR7, CCL19, CCL25, CCR6, CCL20, IL-17, TNF-α, IL-2, IL-2R, IL-10, IL-6, TGF-β, claudin-1, and occludin, and normalized to the housekeeping gene GAPDH. Each reaction contained 25 ng of cDNA in a well with 2× SYBR Green PCR Master Mix (Promega, USA) and the appropriate primers. Threshold cycle values were normalized and expressed as relative gene expression.

### 2.13 16S rRNA microbial community analysis

Fecal pellets from the sham and ovx mouse groups were collected on day 45, placed in sterile tubes, and immediately stored at −80°C for the metagenome study. The commercially available QIAamp DNA stool mini kit (QIAGEN) was used to extract metagenomic DNA from the fecal samples following the manufacturer’s instructions. The quality of the isolated metagenomic DNA was assessed using the A260/280 ratio with a NanoDrop. Samples passing QC were used to generate initial amplicons, and the Nextera XT Index kit (Illumina) was used to prepare the NGS libraries. The Illumina MiSeq platform sequenced the QC-passed libraries. Alpha diversity was calculated for each sample, using the number of observed features (ASVs). Microbiota analysis was performed using the QIIME2 pipeline. High-quality clean reads were obtained with Trimmomatic v0.38, employing a sliding window of 20 bp, a minimum length of 100 bp, and this process removed adapter sequences, ambiguous reads (with more than 5% unknown nucleotides “N”), and low-quality sequences (reads with more than 10% of bases below a QV of 25). PE data were converted into single-end reads using FLASH. Chimeric sequences were filtered out using DADA2 after denoising the high-quality clean reads. Taxonomic classification of amplicon sequence variants was performed using a pre-trained classifier based on the SILVA database within q2-feature. The statistical significance of diversity measures was also evaluated in QIIME2.

### 2.14 Analysis of Short Chain Fatty Acids (SCFAs)

High-performance liquid chromatography (HPLC) was used for SCFA analysis following our previous publication (Bhardwaj et al., 2025). Briefly, 1 ml of Milli-Q water and 100 µl of hydrochloric acid (HCl) were added to 300 mg of fecal sample. The sample was thoroughly homogenized with a vortex (2-3 minutes) and left to stand for 20 minutes. It was then centrifuged at 13850 g for 10 minutes at 4°C. After centrifugation, the supernatant was transferred to a 2 ml Eppendorf tube. To this, 600 µl of diethyl ether was added, and the mixture was extracted for another 20 minutes. Following extraction, the sample was centrifuged again at 850 g for 5 minutes at 4°C, and 400 µl of the supernatant was collected. This aliquot was further mixed with 500 µl of sodium hydroxide (NaOH) and extracted for an additional 20 minutes. The sample was then centrifuged at 850 g for 5 minutes. The ether layer was discarded, and 450 µl of the remaining aqueous layer was collected. To this, 300 µl of HCl was added, and the mixture was immediately filtered through a 0.22-micron filter. The prepared sample was analyzed on the HPLC system. HPLC conditions involved an Xterra C18 column (250 mm x 4.6 mm x 3.5 µm); mobile phase, 10 mM sulfuric acid (H2SO4) (isocratic gradient); flow rate, 0.6 ml/min; run time, 11 minutes; and detection wavelength, 210 nm. SCFA levels were determined using standard calibration curves.

### 2.15 Gut permeability assay (Evans Blue Assay)

Mice received an intravenous injection of Evans Blue dye (30 mg/kg body weight) via the tail vein. After 30 minutes, the mice were euthanized, and the large intestine (LI) was removed and stored for 24 hours at 72°C in formamide solution. Additionally, absorbance was measured at 620 nm after collecting the supernatant.

### 2.16 Statistical Analysis

Statistical differences between groups (mice) were analyzed using either paired or unpaired Student’s t-tests as appropriate. All data are expressed as mean ± SEM. Statistical significance was set at p ≤ 0.05 (*p < 0.05, **p < 0.01, ***p < 0.001, ****p < 0.0001) for the specified groups.

## 3.0. Results

### 3.1. ILC3 inhibits differentiation of osteoclasts

To determine whether ILC3s can influence osteoclast differentiation, we first generated ILC3s under *in vitro* conditions. To do this, immune cells were isolated from the spleen and subjected to magnetic separation to select lineage-negative cells using a biotin-labelled antibody cocktail (anti-CD3, CD19, F4/80, NK1.1, FCεRI, Gr-1, CD5, CD11c, TER119). The negatively selected cell population was induced *in vitro* under ILC3-polarizing conditions (IL-2, IL-7, IL-23, and IL-1β). After 24 hours of stimulation, cells were analyzed by flow cytometry. Once ILC3s were generated *in vitro*, we assessed their direct effect on osteoclast differentiation. For this, BMCs were isolated from mice femurs and co-cultured with ILC3s at different cell ratios (BMCs: ILC3; 1:1 and 1:5) in the presence of MCSF (30 ng/mL) and RANKL (60 ng/mL) **(Figure 1A)**. After 4 days, TRAP staining was performed to identify differentiated multinucleated osteoclasts. Interestingly, results showed that ILC3 treatment significantly reduced osteoclast differentiation in a ratio-dependent manner **(Figure 1B)**, as evidenced by the markedly decreased number of TRAP-positive cells (p<0.01) and cells with more than three nuclei (p<0.01) in the ILC3-treated groups compared to the control group **(Figure 1C-D)**. Additionally, area measurement analysis of multinucleated TRAP-positive cells using ImageJ software further confirmed the significant reduction (p<0.0001) in the osteoclast area in ILC3-treated groups **(Figure 1E)**. Collectively, these findings demonstrate the anti-osteoclastogenic potential of ILC3s.

**Figure 1:**
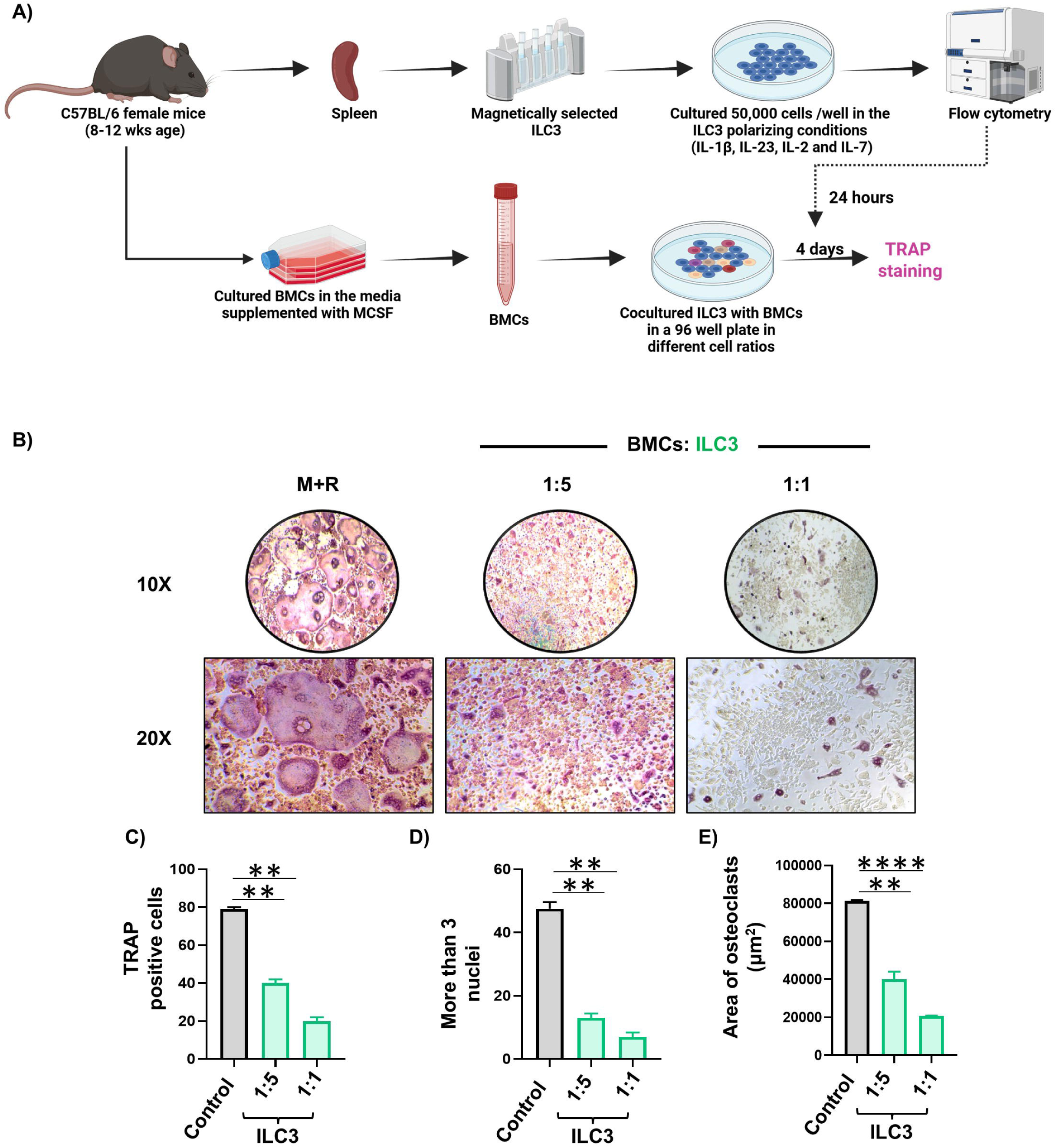
ILC3 inhibits osteoclastogenesis. **(A)** Spleen was harvested and processed for negative selection of ILCs, followed by stimulation with IL-2 (10 ng/ml), IL-7 (10 ng/ml), IL-1β (50 ng/ml), and IL-23 (50 ng/ml), for 24 Hrs. BMCs were harvested from mice and were co-cultured with stimulated ILC3s in the presence of MCSF (30 ng/ml) and RANKL (60 ng/ml). **(B)** ILC3 significantly inhibited the generation of multinucleated osteoclasts in a ratio-dependent manner, as depicted by the pictograph images of the TRAP staining. **(C)** Bar graph representing the number of TRAP^+^ cells**. (D)** Bar graph representing the number of multinucleated cells with more than three nuclei. **(E)** Bar graph representing the area of a multinucleated osteoclast. The results were evaluated using the unpaired Student t-test, and similar results were obtained in two independent experiments. Statistical significance was defined as *p ≤ 0.05, **p < 0.01, *** p ≤ 0.001, **** p ≤ 0.0001 concerning the indicated mouse group.

### 3.2 ILC3 inhibit osteoclastogenesis in an IL-2-dependent manner

ILC3 are reported to majorly secrete two cytokines i.e. IL-17 and IL-22. To determine whether ILC3 inhibits osteoclastogenesis through IL-17 and IL-22, neutralization assays were performed. Anti-IL-17 and anti-IL-22 monoclonal antibodies (10 µg/ml) were added to neutralize these cytokines in cocultures of BMCs and ILC3. Notably, blocking IL-17 and IL-22 did not eliminate the anti-osteoclastogenic effect of ILC3, indicating that ILC3 does not inhibit osteoclastogenesis in an IL-17 or IL-22-dependent manner **(Figures 2A-B)**. Recent studies have shown that ILC3 is the primary producer of IL-2 (Zhou et al., 2019). We too observed that stimulating ILCs with IL-1β and IL-23 increased IL-2 expression **(Figure 2C)**. This suggests that ILC3 might inhibit osteoclastogenesis via IL-2. However, no studies have explored IL-2’s role on osteoclastogenesis. To investigate this, we first examined the gene expression of IL-2 receptor subunits (IL-2Rα, IL-2Rβ, and IL-2γ) on osteoclasts. All subunits of IL-2 receptors were expressed on osteoclasts, with IL-2Rα being the most abundant, implying that IL-2 could influence osteoclastogenesis **(Figure 2D)**. Next, we evaluated the direct effect of IL-2 on osteoclast differentiation. BMCs were cultured with varying concentrations of IL-2 in the presence of M-CSF and RANKL. Interestingly, IL-2 inhibited osteoclastogenesis in a dose-dependent manner, evidenced by a significant reduction in the number of multinucleated cells (more than 3 nuclei) in the IL-2-treated groups **(Figures 2E-F)**. Additionally, the area occupied by osteoclasts was significantly decreased in these groups **(Figure 2G)**. These findings suggest that ILC3 inhibit osteoclastogenesis in an IL-2 dependent manner rather than IL-17 and/or IL-22.

**Figure 2:**
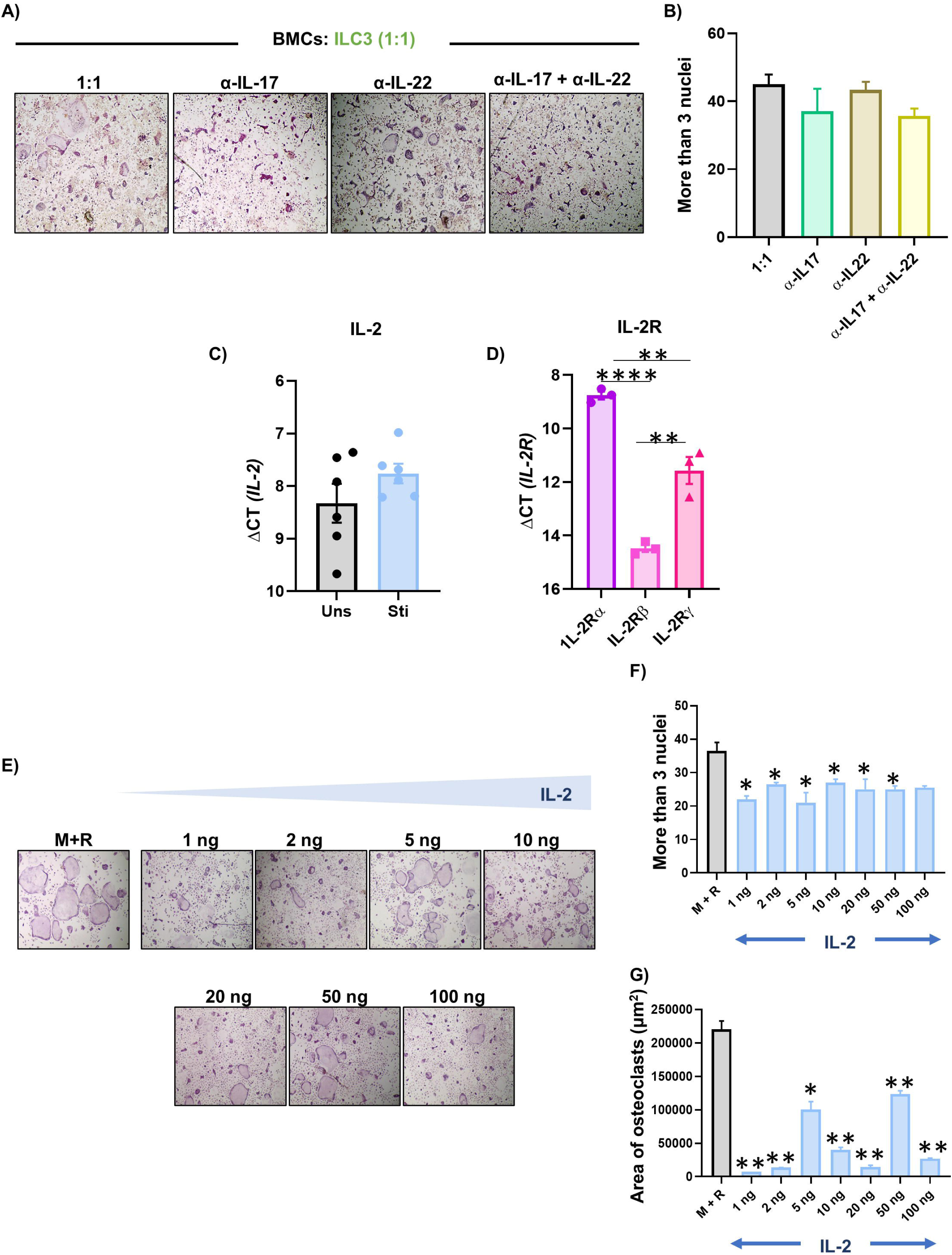
ILC3 inhibits osteoclastogenesis in an IL-17 and IL-22 independent manner. The spleen was harvested and processed for the negative selection of ILCs, followed by stimulation with IL-2 (10 ng/ml), IL-7 (10 ng/ml), IL-1β (50 ng/ml), and IL-23 (50 ng/ml) for 24 Hours. BMCs were harvested from mice and were co-cultured with stimulated ILC3s in the presence of α-IL-17 and α-IL-22, along with MCSF (30 ng/ml) and RANKL (60 ng/ml). **(A)** α-IL-17 and α-IL-22 are unable to abolish the ILC3-mediated inhibition of osteoclastogenesis, as depicted by the pictograph images of the TRAP staining. **(B)** Bar graph representing the number of multinucleated cells with more than three nuclei. **(C)** Gene expression analysis of IL-2 in unstimulated and stimulated ILC3s. **(D)** Gene expression analysis of different subunits of IL-2 receptor (IL-2R) on osteoclast cells. BMCs were harvested from mice and were cultured in the presence or absence of IL-2 along with MCSF (30 ng/ml) and RANKL (60 ng/ml). **(E)** Pictograph images (10X) representing osteoclastogenesis in the presence or absence of IL-2. **(F)** Bar graph representing the number of multinucleated cells with more than three nuclei. **(G)** Bar graph representing the area of a multinucleated osteoclast. (The results were evaluated using the unpaired Student t-test, and similar results were obtained in two independent experiments. Statistical significance was defined as *p ≤ 0.05, **p < 0.01, *** p ≤ 0.001 concerning the indicated mouse group.

### 3.3 Development of Post-Menopausal Osteoporotic (PMO) Mice Model

Moving forward in our study, we next examined the bone health-modulating potential of ILC3s in *in vivo* conditions. To do this, we randomly assigned female C57BL/6 mice into two groups: sham (ovaries intact) and ovx (ovaries bilaterally removed) **(Figure 3A)**. To confirm the successful development of the PMO mouse model, we analyzed the effects of estrogen deficiency on bone resorption and bone histomorphometry parameters. During the 45 days of osteoporosis development, the body weight of mice from both groups was measured at regular intervals (days 1, 15, 30, and 45). No significant changes in body weight were observed at these time points in either sham or ovx mice **(Supplementary Figure 1A)**.

**Figure 3:**
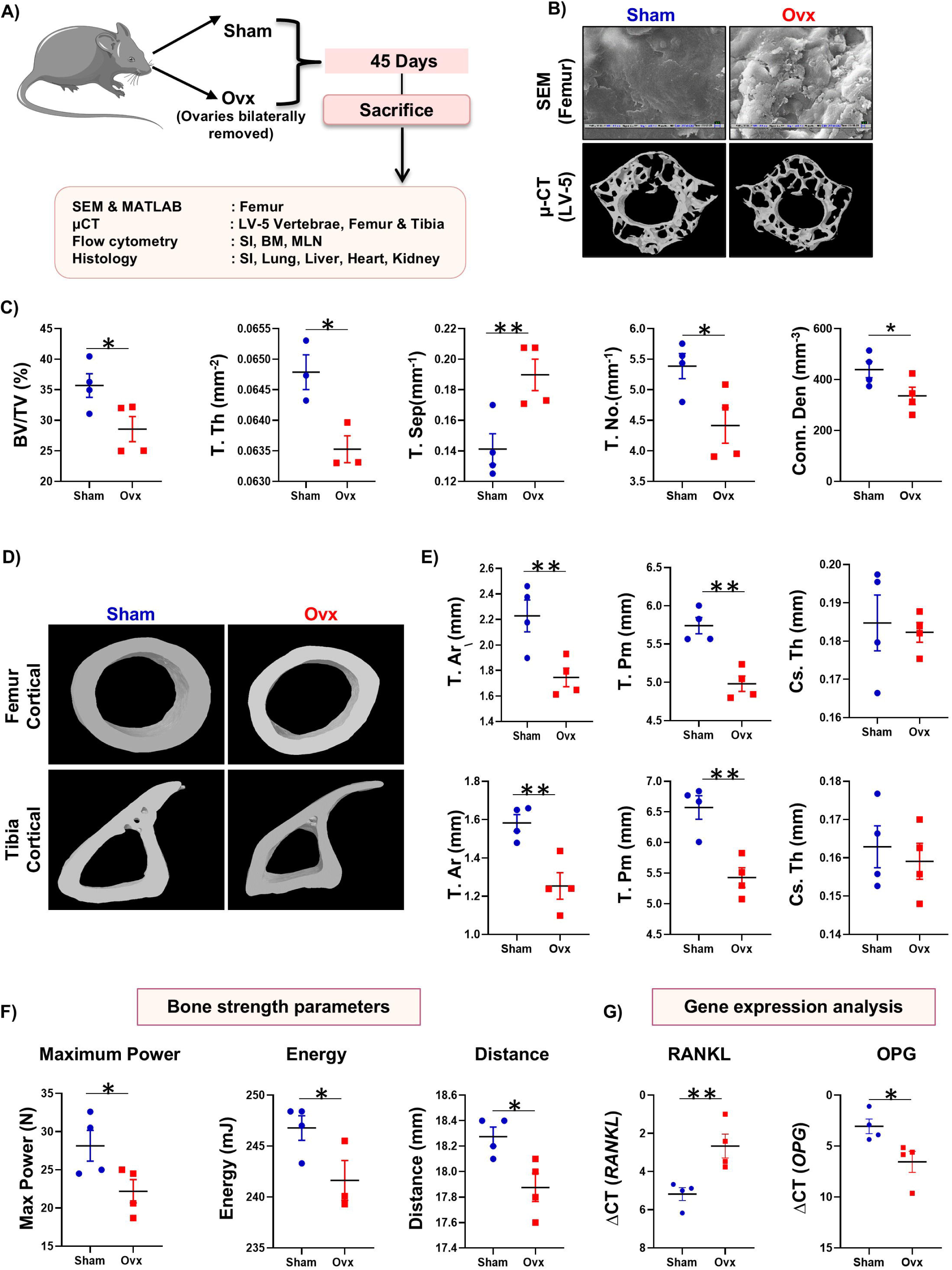
Development of Post-menopausal osteoporotic (PMO) mice model: **(A)** Experiment layout and work plan for *in vivo* study. Mice were divided into two groups, viz. Sham and Ovx in which ovariectomy was performed. **(B)** Scanning electron microscopy (SEM) images of femur cortical bone and 3D µ-CT reconstruction images of lumbar vertebrae (LV)-5 bone in the sham and ovx group. **(C)** Graphical representation of different histomorphometric indices of LV-5 trabecular region of the sham and ovx groups. **(D)** 3D µ-CT reconstruction images of femur and tibia cortical bones in the sham and ovx groups. **(E)** Graphical representation of different histomorphometric indices of femur and tibia cortical regions of the sham and ovx groups. **(F)** Three-point bending test of femur diaphysis representing max power (N), energy (mJ), and distance (mm). **(G)** Gene expression analysis of receptor activator of nuclear factor-κB ligand (RANKL) and osteoprotegerin (OPG) in bone marrow cells. BV/TV: bone volume to total volume; Tb. Th: bone trabecular thickness (Tb. Th); Tb. Sep: trabecular separation; T. No: trabecular number; Conn.Den: connectivity density, T. Ar: total cross-sectional area, T. Pm: total cross-sectional perimeter (T. Pm); Cs. Th: cross-sectional thickness. The results were evaluated using the unpaired Student t-test. Values are expressed as the mean ± SEM (n = 4), and similar results were obtained in two independent experiments. Statistical significance was defined as *p ≤ 0.05, **p < 0.01, *** p ≤ 0.001 concerning the indicated mouse group.

To further evaluate how ovariectomy affects organ weight, the spleen, SI, and colon were harvested, and their weights were measured. A significant increase in the weights of these organs (spleen, SI, and colon) was observed in the ovx group, indicating that ovariectomy promotes adipogenesis in various organs, as also reported in earlier studies (Gavin et al., 2018) **(Supplementary Figure 1B-D)**. Additionally, the ovariectomized mice showed a significant increase in spleen length, suggesting inflammation **(Supplementary Figure 1E-F)**. SEM analysis of the cortical region of femoral bone revealed more resorption pits, or lacunae, in the ovx group compared to sham-operated mice **(Figure 3B)**. Furthermore, µ-CT was used to assess bone microarchitecture, along with histomorphometric analysis of LV-5. The µ-CT data showed that, compared to sham mice, the PMO model mice exhibited significant deterioration in the microarchitecture of LV-5 bones (**Figure 3B)**. Quantitative analysis of LV-5 trabecular bone, which summarizes histomorphometry indices from 3D images of bone microarchitecture, demonstrated significant decreases in BV/TV (p<0.05), T. Th (p<0.05), T. No (p<0.05), and Conn. Den (p<0.05), along with an increase in T. Sep (p<0.01) in the ovx group compared to the sham group **(Figure 3C)**. The µ-CT data further indicated notable impairment in the cortical bones of the femur and tibia in the ovx groups, supported by significant reductions in T.Ar (p<0.05) and T.Pm (p<0.01) compared to sham mice **(Figures 3D-E)**. Additionally, the bone loss observed in the ovx group was further confirmed by the 3-point bending test, which showed significant reductions in maximum force (force the bone can withstand before fracturing) and energy absorption before failure, deformation, and displacement **(Figure 3F)**. The ovx group also exhibited a significant increase in RANKL gene expression (p<0.01) and a decrease in osteoprotegerin (OPG) expression (p<0.05) in the bone marrow cells compared to the sham group **(Figure 3G)**. Overall, these results confirm the successful development of a preclinical mouse model of osteoporosis.

### 3.4. ILC3s play an important role in ameliorating ovariectomy-induced bone loss

To investigate whether our *in vitro* findings of ILC3 could be targeted for new therapeutic strategies in bone loss conditions, we next examined the immunoprotective role of ILC3 in regulating bone health using the PMO mice model. To do this, BM, the main site of osteoclast formation, along with mesenteric lymph nodes (MLN) and small intestine (SI), were harvested from both groups. Flow cytometric analysis for ILC3 was performed according to the gating strategy shown in **Figure 4A**. Notably, compared to the sham group, the ovx group showed a significant decrease in ILC3 population in the BM (p < 0.05) **(Figures 4B-D)**, indicating an important role for ILC3 in maintaining bone health. Conversely, the ILC3 population was significantly increased in the MLN (p<0.05) and SI (p<0.05) in the ovx group **(Figures 4B-D).** While the proportion of ILC3 cells were changed, the mean fluorescence intensity (MFI) of RORγT was higher in all tissues of ovx mice, including BM (p < 0.05), MLN (p < 0.05), and SI (p < 0.05) (Figure 4E). t-SNE plots further confirmed reduced RORγT expression in CD3-negative cells in the BM and increased expression in the MLN and SI **(Figure 4C)**. Additionally, the ILC3 population in the BM showed a significant positive correlation with femoral trabecular BMD, while ILC3s in the MLN and SI exhibited a negative correlation with BMD. Altogether, these results highlight an important role of ILC3s in the pathophysiology of PMO.

**Figure 4:**
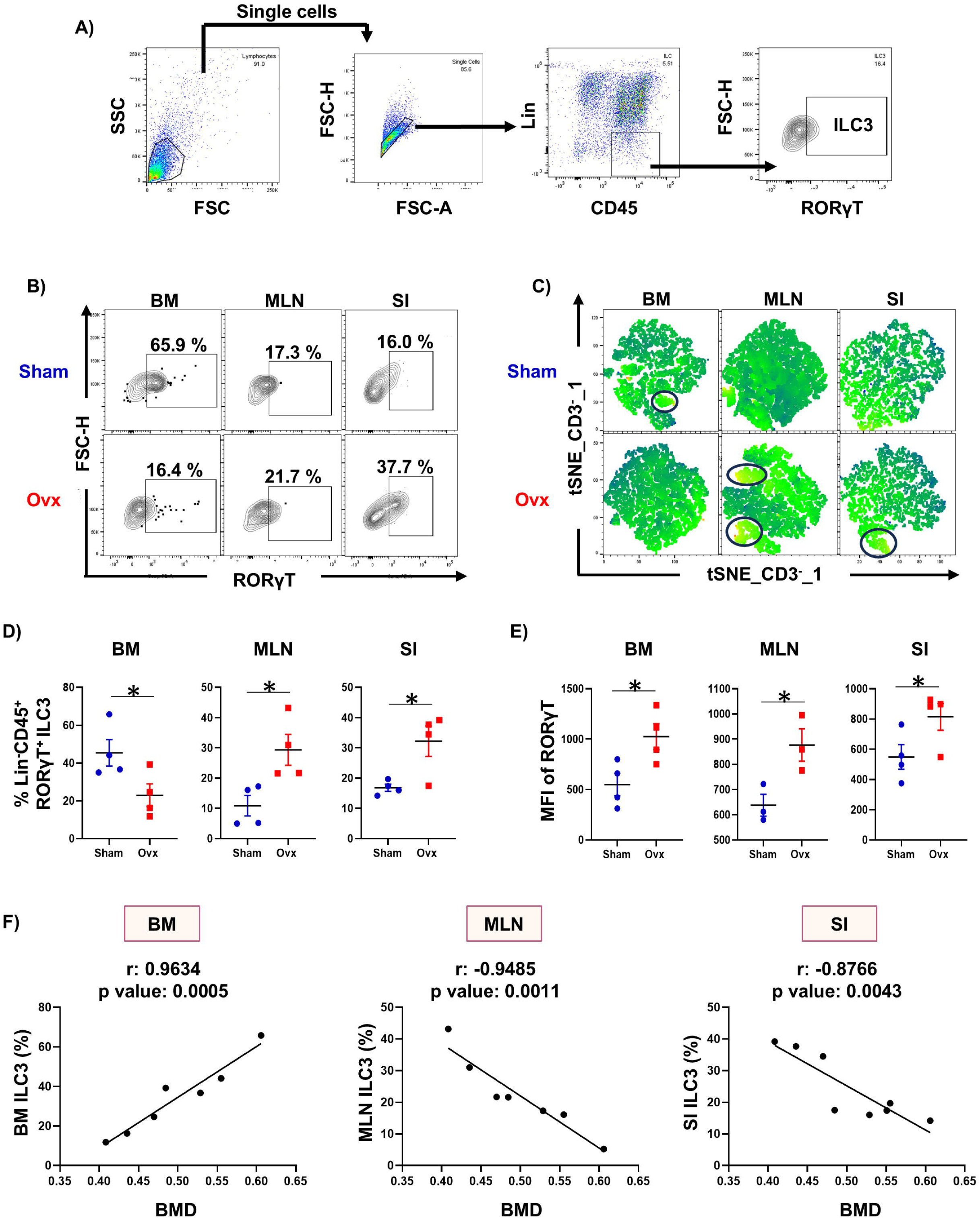
PMO results in modulation of innate lymphoid cell (ILC)-3 *in vivo*. **(A)** Gating strategy adopted for characterization of the ILC3 population. Cells from the bone marrow (BM), mesenteric lymph nodes (MLN), and lamina propria of the small intestine (SI) of sham and ovx groups were harvested and analyzed by flow cytometry for the percentage of ILC3. **(B)** Contour plot depicting percentages of ILC3 (Lin^−^CD45^+^RORγT^+^) in BM, MLN, and SI of sham and ovx groups. **(C)** t-SNE plots representing the RORγT^+^ cells in the CD3^−^ population. **(D).** Bar graph representing percentages of ILC3 in sham and ovx groups. **(E)** Bar graph representing the mean fluorescence intensity (MFI) of RORγT in the sham and ovx groups. **(F)** Correlation graphs depicting the correlation of ILC3 from the BM, MLN, and SI with the bone mineral density (BMD). The results were evaluated using the unpaired Student t-test. Values are expressed as mean ± SEM (n=4), and similar results were obtained in two independent experiments. Statistical significance was defined as *p ≤ 0.05, **p < 0.01, *** p ≤ 0.001 concerning the indicated mouse group.

### 3.5. ILC3s are dynamically altered during the progression of postmenopausal osteoporosis

To determine the role of ILC3 in PMO progression, we analyzed the frequencies of ILC3 in the BM at different time points (days 15, 30, and 45). Notably, even on day 15 post-ovariectomy, ILC3 numbers were significantly reduced in the BM of ovx mice (p < 0.05) **(Figure 5A-B)**. Strikingly, by day 30 post-ovariectomy—which corresponds to the osteopenic stage—the ILC3 population was drastically decreased in the BM (p < 0.05**) (Figure 5A-B)**. Similar results were observed at day 45 post-ovariectomy—indicating osteoporotic conditions—with ILC3s decreased in the BM (p < 0.05) compared to sham mice **(Figure 5A-B)**. Additionally, chemokines and their ligands that regulate ILC3 migration (CCR7-p < 0.05, CCL19-p < 0.001, CCL25-p < 0.01, CCR6-p < 0.01, CCL20-p < 0.01) were significantly reduced in the BM of ovx mice **(Figure 5D)**. Overall, our results highlight that postmenopausal osteoporotic conditions promotes downregulation of chemokine and ligand expression, which further impairs ILC3 migration to the BM, leading to a decreased frequency of ILC3s in the BM of the ovx mice.

**Figure 5:**
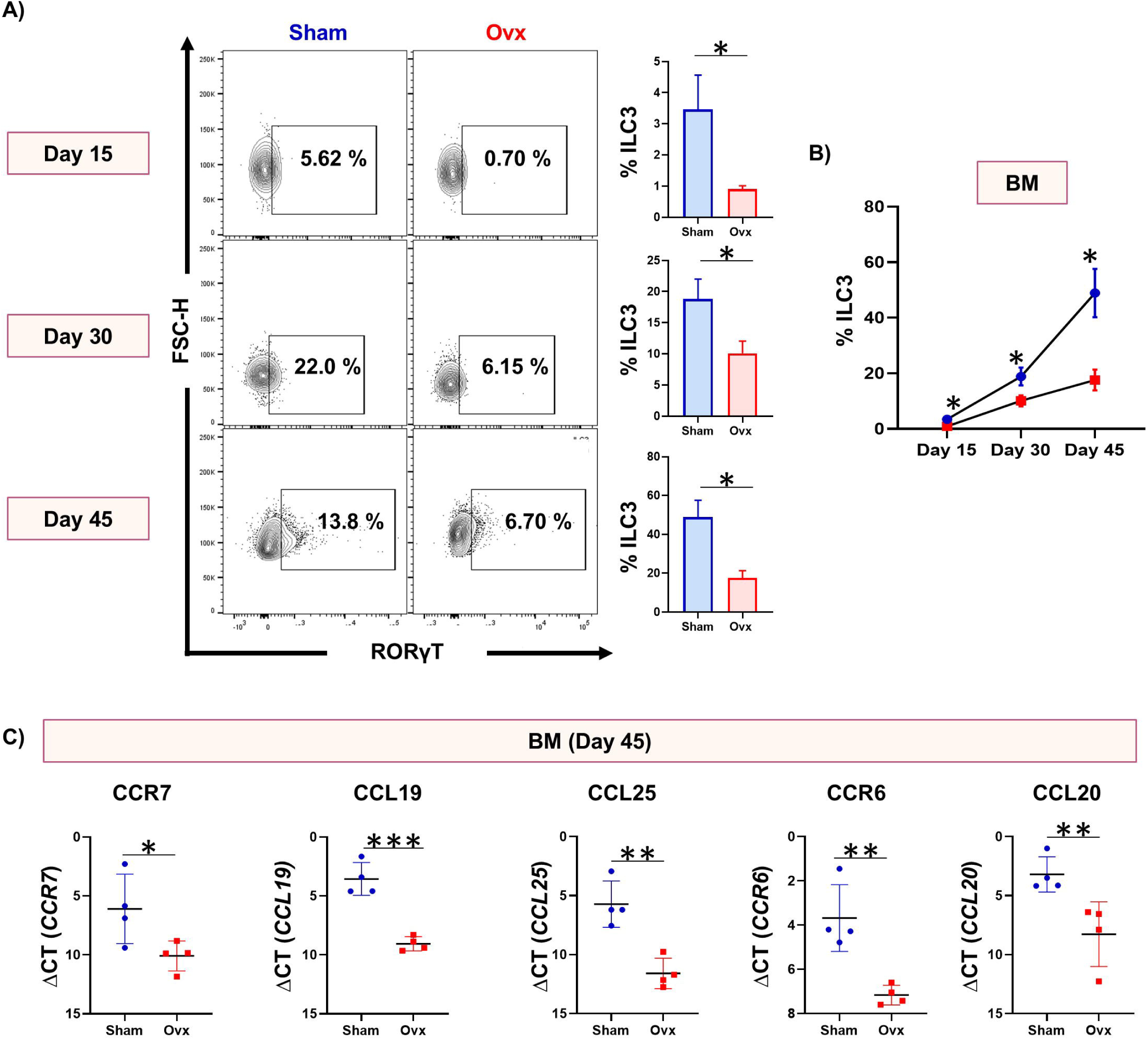
ILC3s are dynamically altered during progression of osteoporosis: **(A-B)** Contour plots, bar graphs, and line plot representing the ILC3 population in bone marrow (BM) of sham and Ovx mice at day 15, 30, and 45. **(C)** Gene expression analysis of different chemokines and their ligands in the BM of sham and ovx groups. CCL: C-C motif chemokine ligand; CCR: C-C chemokine receptor. The results were evaluated using the unpaired Student t-test. Values are expressed as the mean ± SEM (n = 4), and similar results were obtained in two independent experiments. Statistical significance was defined as *p ≤ 0.05, **p < 0.01, *** p ≤ 0.001 concerning the indicated mouse group.

### 3.6 Dysbiosis in PMO promotes the differentiation of IL-17-producing Nkp46^−^ ILC3

Previous studies have shown that increased gut permeability and dysbiosis disrupt ILC3 homeostasis (Hu et al., 2023). To determine whether PMO is linked to disturbed ILC3 balance, we first conducted gut permeability tests and gut microbiome analyses in the sham and ovx groups. Notably, significant histological changes were observed in the small intestine (SI) of the ovx group, aligning with previous findings that ovariectomy induces chronic intestinal inflammation **(Figure 6A)**. We also found that gut permeability was significantly higher in the ovx group **(Figure 6B)**. Additionally, in the ovx group, the levels of tight junction proteins, claudin-1 and occludin, in the large intestine were markedly reduced, suggesting that PMO is associated with a leaky gut and dysbiosis **(Figure 6C-D)**. We thus next examined the gut microbiome composition in fecal samples from both groups. Dominant bacterial phyla included Firmicutes, Bacteroidetes, Patescibacteria, Proteobacteria, and Actinobacteria. In the ovx group, the abundance of Firmicutes and Bacteroidetes decreased, while Desulfobacterota—linked to increased gut permeability—was elevated **(Figure 6E-F)**. Differences in alpha diversity were also observed between the groups **(Figure 6G)**. At the family level, families involved in SCFA production, such as Lactobacillaceae, Bacteriaceae, and Clostridia_UCG-014 (highlighted in green), were reduced in the ovx group. Conversely, families associated with inflammation and leaky gut, like Desulfovibrionaceae (highlighted in red), were increased in ovx mice **(Figure 6H)**. Similar trends were seen at the genus level, with SCFA-producing genera such as Lactobacillus, Bacteroides, and Clostridia_UCG-014 being decreased in the ovx group **(Figure 7A)**. Krona charts further confirmed that the class Clostridia, a major SCFA producing group, was reduced in ovx mice **(Figure 7B)**. These findings suggest that dysbiosis in ovx mice correlates with reduced SCFA production. To validate our microbiome data, we performed targeted metabolomics on fecal samples, which showed significantly lower levels of SCFAs in the ovx group compared to the sham group, supporting our above metagenomic results **(Figure 7C)**.

**Figure 6:**
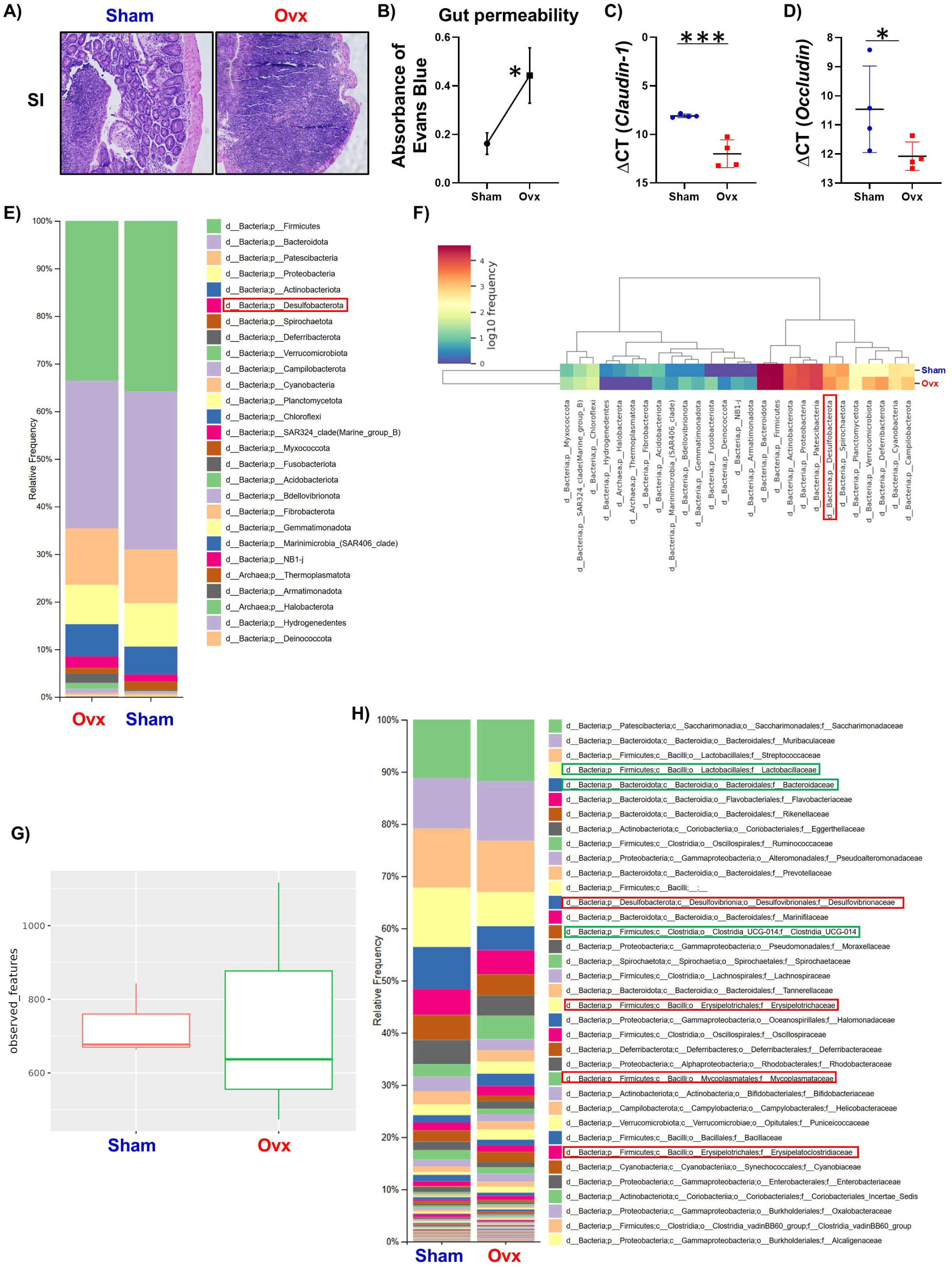
PMO is associated with loss of gut-barrier integrity and gut dysbiosis in Ovx mice: **(A)** H&E staining of small intestine tissues. **(B)** Measurement of gut integrity in sham and Ovx-operated mice. **(C-D)** Relative gene expression of tight junctional proteins occludin and claudin in the large intestine. **(E)** Stacked bar chart representing the relative abundance of each phylum within the sham and ovx groups. **(F)** Heatmap denoting the dominant phylum in each group. (**G)** Bar graph representing alpha diversity in each group. **(H)** Stacked bar chart representing the relative abundance of each family within the sham and ovx groups. Values are expressed as the mean ± SEM (n = 4), and similar results were obtained in two independent experiments. Statistical significance was defined as *p ≤ 0.05, **p < 0.01, *** p ≤ 0.001 concerning the indicated mouse group.

**Figure 7:**
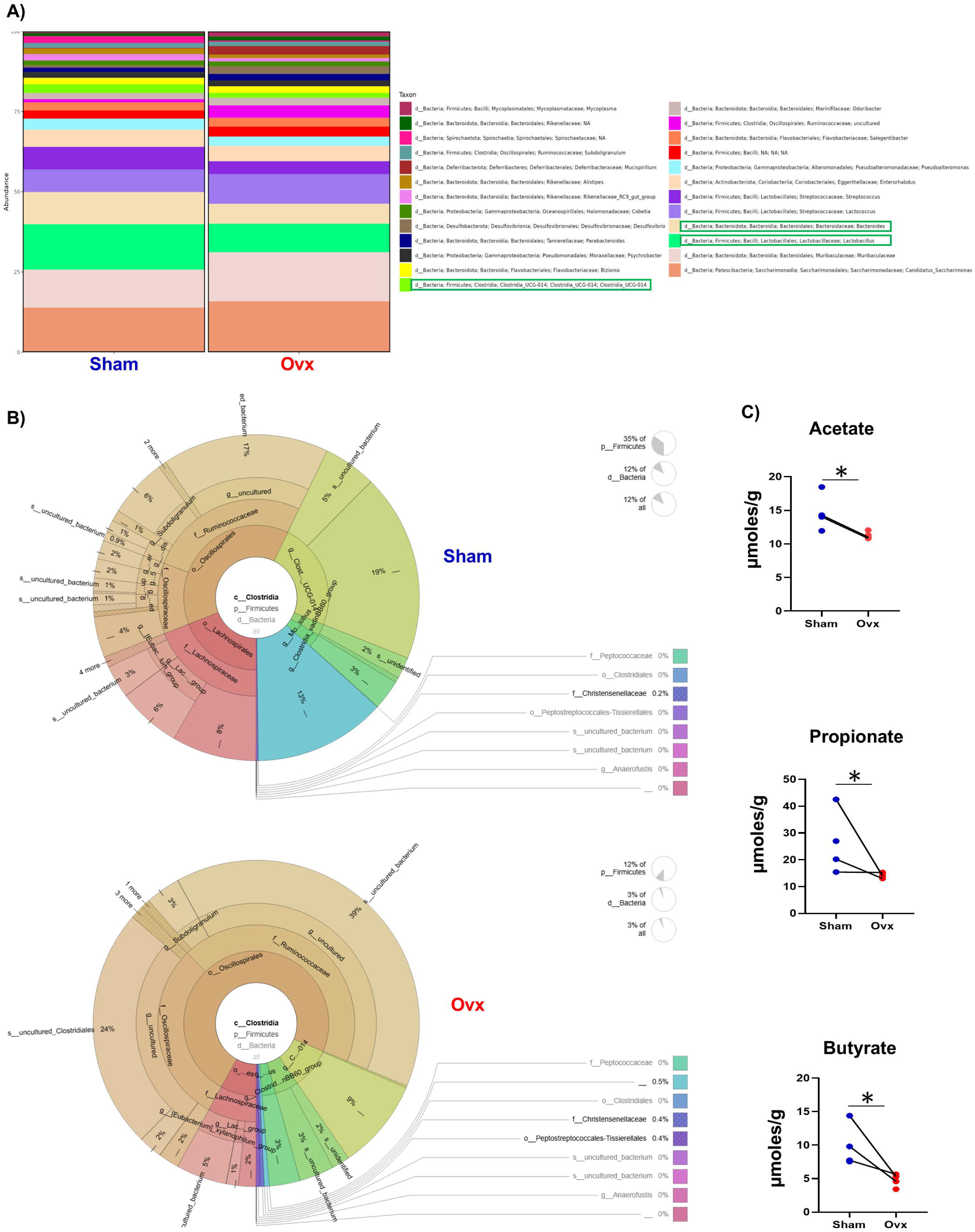
Dysbiosis results in short-chain fatty acids (SCFAs) depletion in Ovx mice: **(A)** Stacked bar chart representing the relative abundance of each genus within the sham and ovx groups. **(B)** Krona charts representing the taxonomic distribution of Clostridia in sham and ovx groups. **(C)** SCFAs (acetate, propionate and butyrate) analysis in the fecal samples of sham and ovx groups. The results were evaluated using the unpaired Student t-test. Values are expressed as the mean ± SEM (n = 4), and similar results were obtained in two independent experiments. Statistical significance was defined as *p ≤ 0.05, **p < 0.01, *** p ≤ 0.001 concerning the indicated mouse group.

Earlier studies indicated that SCFAs help maintain ILC3 homeostasis (Hu et al., 2023). Therefore, we examined the effect of SCFA depletion on ILC3s *in vivo*. ILC3s are divided into two subsets: IL-17-producing and IL-22-producing. SCFAs appear to inhibit the development of IL-17-producing ILC3s while promoting IL-22-producing ILC3s. Next, we determined the percentage of IL-17-producing ILC3s (CD3^−^RORγT^+^IL-17^+^) in BM, MLN, and SI. Our flow cytometry data showed that the ovx group had a significant increase in the percentage of CD3^−^RORγT^+^IL-17^+^ cells across all tissues (p<0.05), along with increased MFI of IL-17 in MLN compared to controls **(Figure 8A-B)**. Additionally, we observed significantly higher gene expression levels of IL-17 in both SI (p<0.001) and BM (p<0.001) **(Figures 8-D)**. Serum IL-17 levels were also significantly elevated (p<0.0001) **(Figure 8E)**. We next analyzed the percentage of IL-22-producing ILC3s in BM, MLN, and SI by flow cytometry. Remarkably, CD3^−^RORγT^+^IL-22^+^ cells were significantly reduced in the gut tissue MLN of ovx mice **(Figure 8F)**. Since IL-17 is mainly produced by NKp46^−^ ILC3s, while IL-22 is mainly produced by NKp46^+^ ILC3s, we further evaluated these cells in all tissues. We found that NKp46^−^ ILC3s were significantly increased (p<0.05), whereas NKp46^+^ILC3s (p<0.05) were decreased in the BM, MLN, and SI of the ovx group compared to sham controls **(Supplementary Figure 2A & Figures 8G-H)**. Interestingly, we further observed that ILC3 from ovx mice showed increased expression of T-bet compared to the control in BM (p<0.05), MLN (p<0.05), and SI (p<0.01) **(Supplementary Figure 2B & Figure 8I-J)**. T-bet^+^ ILC3s have been reported to increase in various inflammatory diseases, including bone disorders like ankylosing spondylitis, and are linked to higher disease activity (Liu et al., 2025). Altogether, our results clearly suggest that SCFA deficiency could lead to enhanced levels of inflammatory IL-17-producing NKp46^−^ ILC3s thereby leading to enhanced bone loss in PMO.

**Figure 8:**
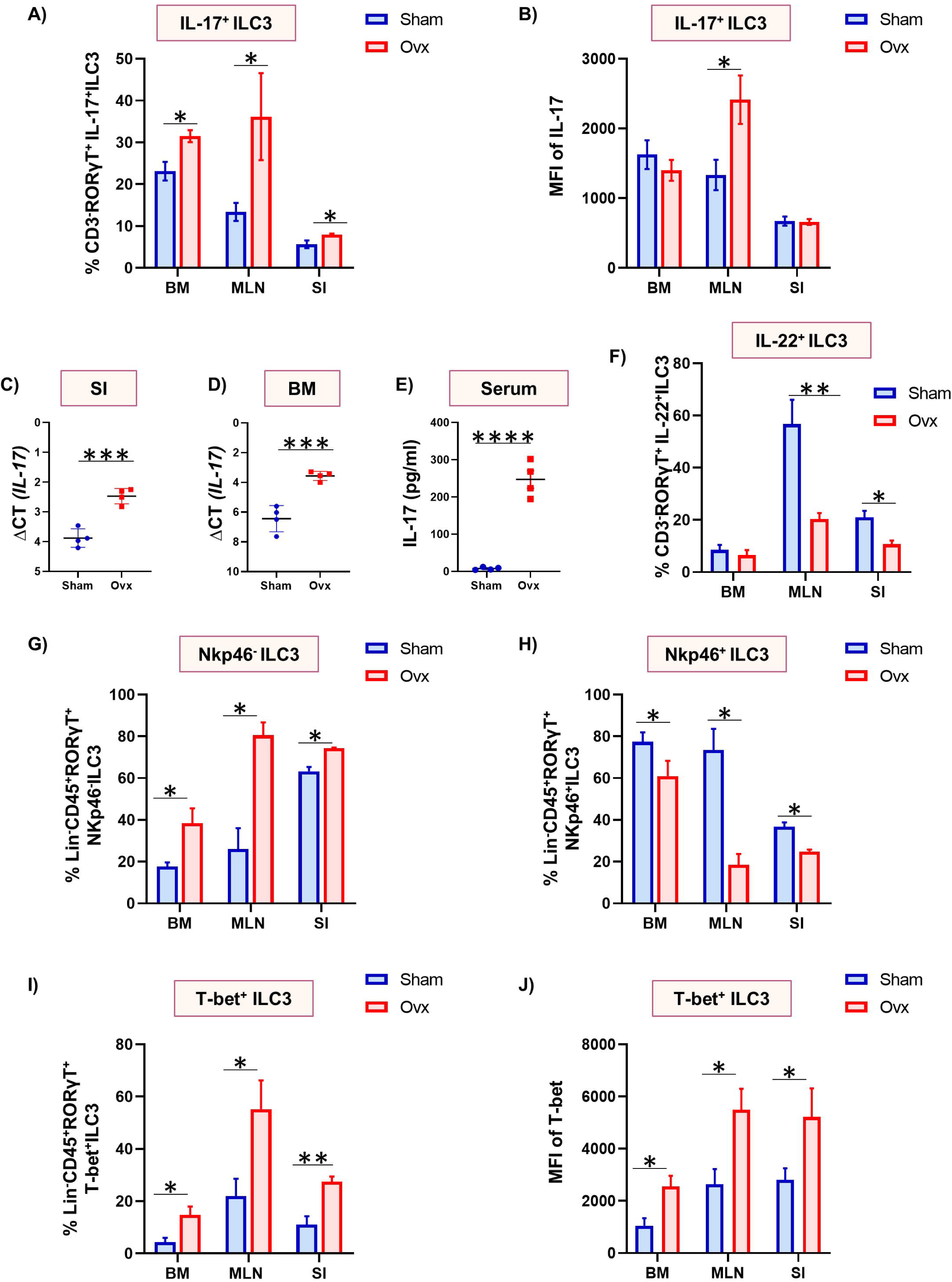
PMO modulates the IL-17^+^ and IL-22^+^ ILC3 populations. Analysis of IL-17 cytokine via flow cytometry in CD3^−^RORγT^+^ ILC3 cells in BM, mesenteric lymph node (MLN), and SI of sham and ovx groups. **(A-B)** Bar graphs representing the CD3^−^RORγT^+^IL-17^+^ ILC3 population and mean fluorescence intensity (MFI) of IL-17 in sham and ovx groups. **(C-D)** Gene expression analysis of IL-17 in the small intestine (SI) and bone marrow (BM) of sham and ovx groups. **(E)** The level of IL-17 cytokine in serum. **(F)** Bar graphs representing the CD3^−^RORγT^+^IL-22^+^ ILC3 population in sham and ovx groups. (**G-H)** Bar graphs representing the NKp46^−^ ILC3 (Lin-CD45^+^RORγT^+^NKp46^−^) and NKp46^+^ ILC3 (Lin-CD45^+^RORγT^+^NKp46^+^) population in the BM, MLN, and SI of sham and ovx groups. (**I)** Bar graph representing the T-bet^+^ ILC3 (Lin-CD45^+^RORγT^+^T-bet^+^) population in BM, MLN, and SI of sham and ovx groups. **(J)** Mean fluorescence intensity (MFI) of T-bet in ILC3 cell population. The results were evaluated using the unpaired Student t-test. Values are expressed as the mean ± SEM (n = 4), and similar results were obtained in two independent experiments. Statistical significance was defined as *p ≤ 0.05, **p < 0.01, *** p ≤ 0.001 concerning the indicated mouse group.

### 3.7 Cytokine imbalance results in compromised ILC3 in ovx mice

Dysbiosis and leaky gut increased levels of inflammatory cytokines IL-6 and TNF-α, which are responsible for the reduced secretion of IL-22 (Powell et al., 2015; Xiong et al., 2025) and increased secretion of IL-17 from ILC3. Therefore, we further analyzed the expression of these cytokines in BM and SI. Gene expression of IL-6 and TNF-α was significantly elevated in both BM and SI **(Figure 9A-B)**. However, the expression of anti-inflammatory cytokines TGF-β and IL-10 was decreased in both BM and SI in the ovx group **(Figure 9A-B)**. Overall, the results highlight that cytokine imbalance in ovx mice leads to increased secretion of IL-17 and decreased secretion of IL-22 from ILC3, resulting in increased inflammation and gut permeability. Since ILC3 is the primary producer of IL-2; therefore, we further examined IL-2 expression in BM and SI. It was observed that IL-2 levels were significantly decreased in the BM and SI of ovx mice **(Figure 9C)**. Conclusively, our results clearly establish the osteoprotective role of ILC3 in ovx mice.

**Figure 9:**
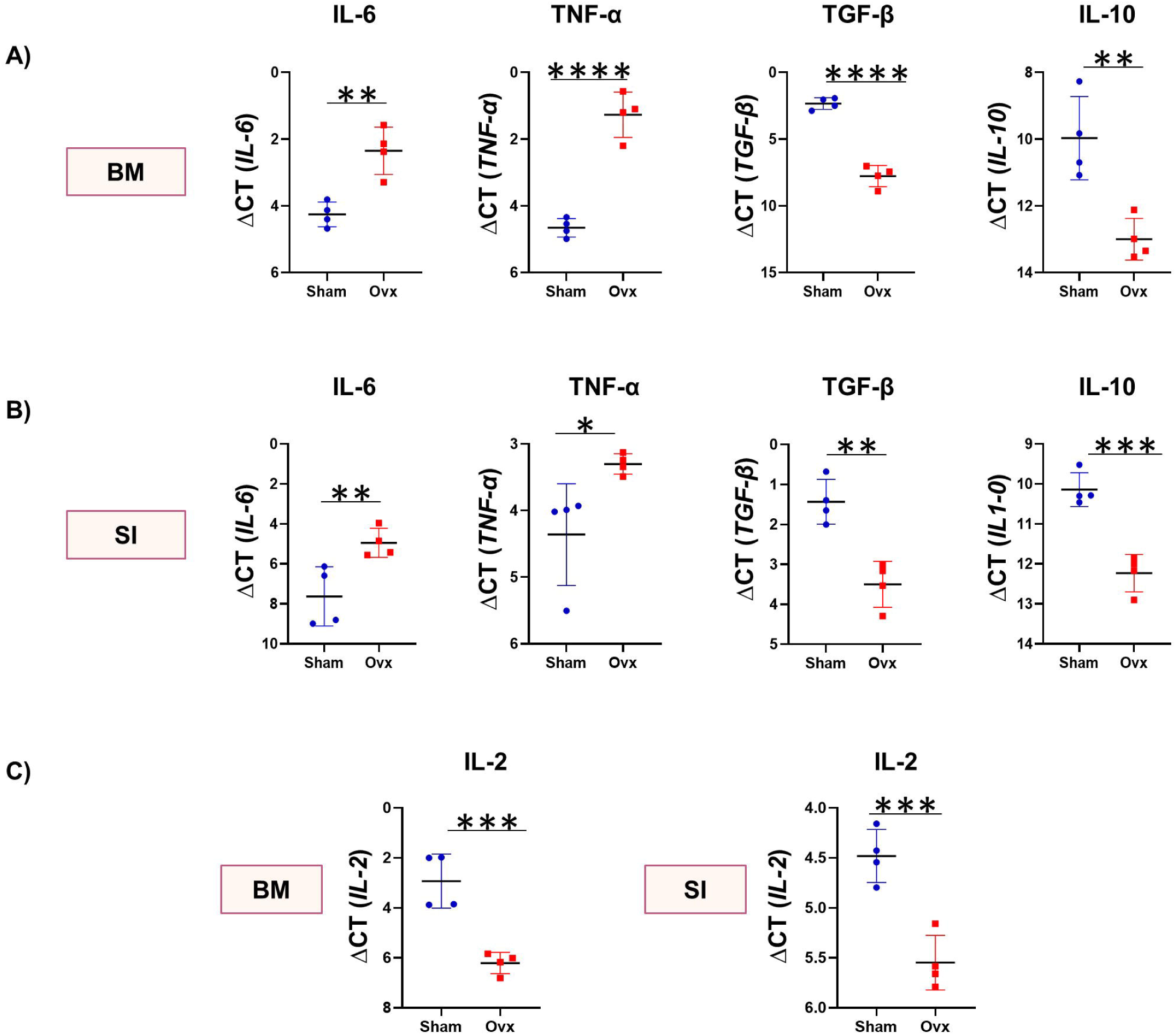
PMO perturbs the cytokine balance in bone marrow and intestine. **(A-B)** Gene expression analysis of different cytokines in the sham and ovx groups. **(C)** Gene expression analysis of IL-2 in the sham and ovx groups. cells in the sham and ovx groups. IL: interleukin; TNF: tumour necrosis factor; TGF: transforming growth factor. The results were evaluated using the unpaired Student t-test. Values are expressed as the mean ± SEM (n = 4), and similar results were obtained in two independent experiments. Statistical significance was defined as *p ≤ 0.05, **p < 0.01, *** p ≤ 0.001 concerning the indicated mouse group.

## 4.0. Discussion

The significance of the immune system in maintaining bone health under both normal and osteoporotic conditions has been highlighted by research from our lab as well as others. These studies led to the creation of the new field of “immunoporosis” (by our group), which specifically emphasizes the role of the immune system in the development of osteoporosis (Pol et al., 2023; Sapra, Azam, et al., 2021; Saxena et al., 2021; Srivastava et al., 2018; Srivastava & Sapra, 2022; Zhang et al., 2022).

ILCs are newly identified immune cells with multiple functions; they have the potential to contribute to early immune responses following infection due to their rapid secretion of immunoregulatory cytokines. It is well known that, despite lacking antigen specificity, they can sense changes in the local environment caused by pathogenic invasion or tissue injury and thus influence subsequent lymphocyte responses (Walker et al., 2013). These ILCs are associated with several inflammatory diseases that lead to bone loss. The role of ILCs in bone disorders such as rheumatoid arthritis (RA), periodontitis, ankylosing spondylitis, and osteoporosis is currently being studied. In a post-menopausal mouse model of osteoporotic bone loss, the adoptive transfer of ILC2 was found to reverse bone loss. The synthesis of cytokines IL-4/IL-13 and activation of STAT6 in osteoclast (OC) progenitors were linked to ILC2’s anti-osteoclastogenic and anti-osteoporotic effects. These findings suggest that ILC2s may play a role in maintaining bone homeostasis (Omata et al., 2020). Since ILCs mirror T helper cells and the Th17/Treg balance (Dar, Lone, et al., 2018; Dar, Pal, et al., 2018; Dar, Shukla, et al., 2018; Dar, Singh, et al., 2018; Sapra, Dar, et al., 2021; Sapra et al., 2022) is critical for bone health, ILC3 may also have bone-regulatory functions. Some conflicting studies regarding RA explore the role of ILC3 in pre-RA and early RA patients. Studies have shown that ILC3 plays an immune regulatory role in humans and mice (Bugaut & Aractingi, 2021b; Li et al., 2021). However, it is difficult to distinguish whether ILC3s contribute to bone maintenance or promote inflammation and bone loss. Additionally, ILC3s secrete cytokines like IL-17, which encourage osteoclast differentiation. Conversely, ILCs also produce IL-22, which helps maintain gut integrity in osteoporosis (Pan et al., 2021; Withers & Hepworth, 2017). Therefore, ILC3s can act as a double-edged sword of the immune system. Given that ILC3s are associated with inflammation and probably indirectly contribute to bone loss in inflammatory diseases, their direct impact on bone remains unproven.

ILCs exhibit plasticity in their nature (Bal et al., 2020); therefore, we chose this particular trait for the generation of ILC3s in our study. We developed and stimulated ILC3 with IL-2, IL-7, IL-1β, and IL-23 for 24 hours. As our research progressed, we became more interested in ILC3’s potential to regulate RANKL-induced osteoclastogenesis in an *in vitro* setting. A remarkable finding was that ILC3s blocked osteoclast differentiation from BMCs in a ratio-dependent manner. Consequently, our study for the first time established the direct role of ILC3s in inhibiting osteoclastogenesis. ILC3s mainly produce cytokines IL-17 and IL-22, both of which are osteoclastogenic cytokines. This raises the question of how ILC3s inhibit bone loss. To determine whether ILC3s suppress osteoclast differentiation in an IL-17 and IL-22-dependent manner, we cocultured ILC3s with BMCs in the presence of α-IL-17 and α-IL-22 and observed that these cytokines do not influence the anti-osteoclastogenic effect of ILC3s. A study has reported that ILC3s also produce IL-2, with the majority of IL-2 coming from ILC3s rather than T cells (Zhou et al., 2019). Therefore, ILC3s may inhibit osteoclastogenesis via IL-2. However, no study has yet evaluated the role of IL-2 in osteoclastogenesis. Hence, we examined whether IL-2 can inhibit osteoclastogenesis. Interestingly, it was observed that osteoclasts express the IL-2 receptor, particularly IL-2Rα, and that IL-2 inhibits osteoclast differentiation in a concentration-dependent manner. Thus, ILC3s could inhibit osteoclastogenesis through an IL-2-dependent mechanism.

Next, we aimed to examine the effect of ILC3s on bone health *in vivo* using the PMO mice model. Interestingly, we found a significant reduction in the overall ILC3 population in the BM of osteoporotic mice, supporting our *in vitro* findings. Conversely, the ILC3 population increased in the MLN and SI. The temporal kinetics study showed that ILC3 levels started to decline in the BM of ovx mice just 15 days after ovariectomy. It has also been observed that chemokines (CCR6 and CCR7) and their ligands (CCL19, CCL20, and CCL25), which guide ILC3 migration from the MLN and intestine, are significantly decreased in the BM. This suggests that the PMO condition may impair the migration of ILC3s to the BM thereby enhancing bone loss.

Previous studies indicated that increased gut permeability and dysbiosis disturb ILC3 homeostasis. To determine whether ILC3 homeostasis is disrupted in ovx mice, we first assessed gut permeability and gut microbiota composition in these mice. Our results showed a significant increase in gut permeability associated with dysbiosis in ovx mice. Interestingly, we observed that the gut microbiota composition was significantly altered, with genera involved in SCFA production notably decreased. Additionally, targeted metabolomics revealed significantly lower levels of SCFAs in ovx mice compared to sham controls. It is known that SCFAs inhibit the development of IL-17-producing ILC3 and promote IL-22-producing ILC3. IL-17 stimulates osteoclastogenesis (Lacey et al., 2009; Le Goff et al., 2019; Song et al., 2019). Consistent with our findings, several studies have reported IL-17 upregulation in PMO. Deleting the IL-17 receptor prevented ovx-induced bone loss, emphasizing IL-17’s central role in osteoporosis development (Le Goff et al., 2019). Moreover, our *in vivo* data showed increased CD3^−^RORγT^+^IL-17^+^ ILC3 cells in BM, MLN, and SI. Notably, we also found that CD3^−^RORγT^+^IL-22^+^ ILC3 cells, which help maintain gut permeability, were reduced in MLN of PMO mice, underscoring IL-22-producing ILC3’s role in preserving gut integrity and limiting inflammation. Since IL-17 is produced by Nkp46^−^ ILC3, we analyzed this subset in ovx mice. Surprisingly, NKp46^−^ ILC3 levels increased in BM, MLN, and SI, while NKp46^+^ IL-22-producing ILC3 decreased. These findings indicate an increase in IL-17-producing NKp46^−^ ILC3 in ovx mice. Additionally, T-bet expression was elevated in the ILC3 of ovx mice. T-bet^+^ ILC3 have been linked to several inflammatory conditions, including bone diseases like ankylosing spondylitis, and tend to be associated with increased disease activity (Liu et al., 2025). Furthermore, it has been observed that cytokines regulating the secretion of IL-17 and IL-22 from ILC3 are affected in the ovx mice, leading to decreased IL-22 production and increased IL-17 release from ILC3. Additionally, IL-2 secretion is reduced in the ovx mice, as IL-2 is primarily produced by ILC3, so lower IL-2 levels may be linked to decreased ILC3s activity. The decrease in IL-2 expression further promotes osteoclastogenesis, worsening bone loss.

Taken together, our results for the first time report the anti-osteoclastogenic role of ILC3s, which is compromised under PMO conditions. Since ILC3s are innate immune cells involved in activating the adaptive immune system, targeting these cells offers a novel promising and effective therapeutic approach for the treatment and management of PMO.

## Acknowledgment

AB, LS, SPK, CS, and RKS acknowledge the Department of Biotechnology, AIIMS, New Delhi, India, for providing infrastructural facilities. AB and LS thank ICMR for the research fellowship. SPK thanks DBT for the research fellowship. CS thanks CCRH-Aysuh for the research fellowship

## Conflict of Interest Statement

The authors declare no conflicts of interest.

## Data availability

All data generated or analyzed during this study are included in this published article and its supplementary information files. The current study’s data are available from the corresponding author upon reasonable request.

## Funding

This work was financially supported by projects: Department of Biotechnology (BT/PR41958/MED/97/524/2021), ICMR (61/05/2022-IMM/BMS), and ICMR (EMDR/IG/13/2024-01-00842).

## Credit authorship contribution statement

RKS contributed to the conception and design of the study; AB, LS, SPK, and CS contributed to the acquisition and analysis of data; and AB contributed to drafting the text or preparing the figures. PKM provided cytokine data and valuable inputs. All authors reviewed the manuscript. All authors contributed to the article and approved the submitted version.

## Notes

### Competing Interest Statement

The authors have declared no competing interest.

