## Supplementary Figures for "ILC3 inhibits Osteoclastogenesis and ameliorates Inflammatory Bone Loss in Post-menopausal Osteoporosis"

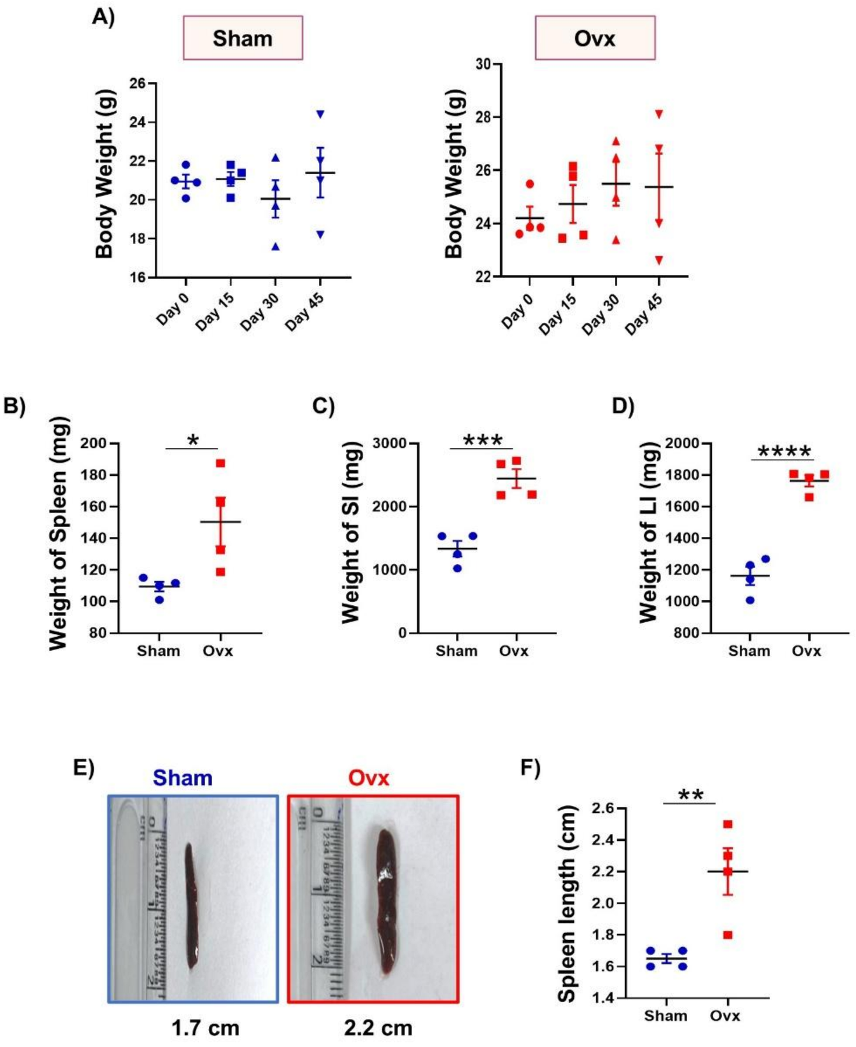


**Supplementary Figure 1. (A)** Body weight of sham and ovx mice at different time points. Effect of ovariectomy on the weight of organs, viz. **(B)** spleen, **(C)** small intestine (SI), and **(D)** large intestine (LI). **(E & F)** Length of spleen in Sham and Ovx mice.

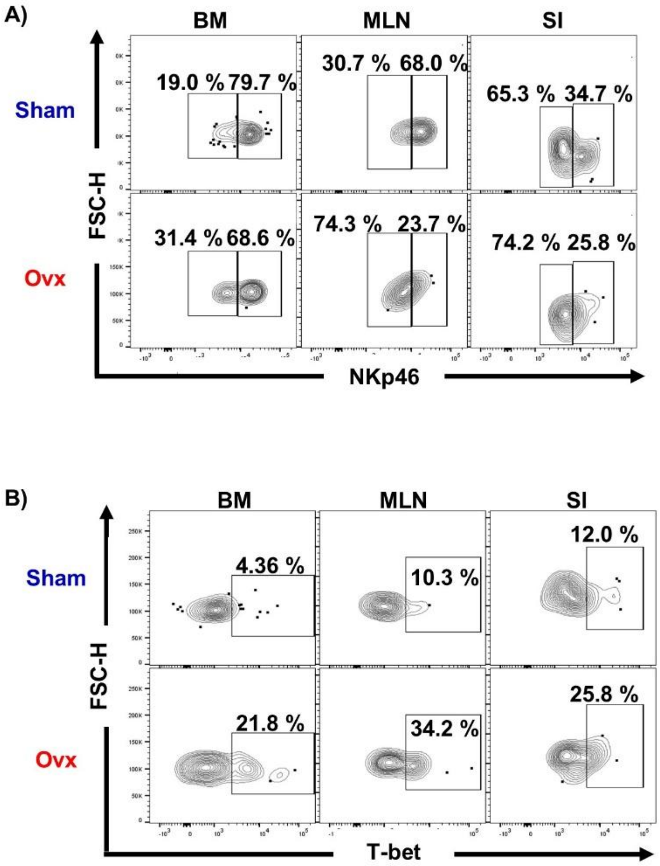


**Supplementary Figure 2: PMO alters the population of ILC3 subsets. (A)** Contour plots representing the NKp46$^-$ ILC3 (Lin-CD45$^+$ROR$\gamma$T$^+$NKp46$^-$) and NKp46$^+$ ILC3 (Lin-CD45$^+$ROR$\gamma$T$^+$NKp46$^+$) population in the BM, MLN, and SI of sham and ovx groups. (**B)** Contour plots representing the T-bet$^+$ ILC3 **(**Lin-CD45$^+$ROR$\gamma$T$^+$T-bet$^+$) population in BM, MLN, and SI of sham and ovx groups.
